# TRPV1 controls innate immunity during *Citrobacter rodentium* enteric infection

**DOI:** 10.1101/2023.07.26.550772

**Authors:** Michael Cremin, Emmy Tay, Valerie T. Ramirez, Kaitlin Murray, Rene K. Nichols, Ingrid Brust-Mascher, Colin Reardon

**Affiliations:** Department of Anatomy, Physiology and Cell Biology, UC Davis School of Veterinary Medicine, UC Davis, Davis, California, USA

## Abstract

Mucosal immunity is critical to host protection from enteric pathogens and must be carefully controlled to prevent immunopathology. Regulation of immune responses can occur through a diverse range of mechanisms including bi-directional communication with the neurons. Among which include specialized sensory neurons that detect noxious stimuli due to the expression of transient receptor potential vanilloid receptor 1 (TRPV1) ion channel and have a significant role in the coordination of host-protective responses to enteric bacterial pathogens. Here we have used the mouse-adapted attaching and effacing pathogen *Citrobacter rodentium* to assess the specific role of the TRPV1 channel in coordinating the host response. TRPV1 knockout (TRPV1^-/-^) mice had a significantly higher *C. rodentium* burden in the distal colon and fecal pellets compared to wild-type (WT) mice. Increased bacterial burden was correlated with significantly increased colonic crypt hyperplasia and proliferating intestinal epithelial cells in TRPV1^-/-^ mice compared to WT. Despite the increased *C. rodentium* burden and histopathology, the recruitment of colonic T cells producing IFNγ, IL-17, or IL-22 was similar between TRPV1^-/-^ and WT mice. In evaluating the innate immune response, we identified that colonic neutrophil recruitment in *C. rodentium* infected TRPV1^-/-^ mice was significantly reduced compared to WT mice; however, this was independent of neutrophil development and maturation within the bone marrow compartment. TRPV1^-/-^ mice were found to have significantly decreased expression of the neutrophil-specific chemokine *Cxcl6* and the adhesion molecules *Icam1* in the distal colon compared to WT mice. Corroborating these findings, a significant reduction in ICAM-1 and VCAM-1, but not MAdCAM-1 protein on the surface of colonic blood endothelial cells from *C. rodentium* infected TRPV1^-/-^ mice compared to WT was observed. These findings demonstrate the critical role of TRPV1 in regulating the host protective responses to enteric bacterial pathogens, and mucosal immune responses.

**Author Summary:** Neuroimmune communications are vital in regulating the immune response to invading pathogens. Here, we show that during a gastrointestinal infection, pain-sensing neuronal fibers can modulate the immune response to recruit phagocytic neutrophils via upregulation of cell adhesion molecules on local blood endothelial cells. This research elucidates a novel impact of the pain-sensing ion channel, TRPV1, on host-pathogen interactions in the gastrointestinal tract as well as a potential methodology for modulating the immune response during enteric infections.

## Introduction

Mucosal host defenses in the intestinal tract are the result of many complex interactions between a diversity of cell types. Host protective immune responses to enteric bacterial pathogens are coordinated by a multitude of factors including the nervous system. The intestinal tract is densely innervated by the highly specialized nociceptive sensory neurons, that can detect noxious stimuli, including danger-associated molecular patterns (e.g. ATP), inflammation, and bacterial products. Detection of these substances followed by the activation of neurons is due, in part, to the expression of the polymodal nociceptor transient receptor potential cation channel subfamily V member 1 (TRPV1) (1, 2). These nociceptive sensory neurons are capable of inducing activation of other neurons in a classical reflex arc that involves the coordinated release of peptidic neurotransmitters (neuropeptides) locally in the intestine (3–5). These neuropeptides, including substance P (SP) and calcitonin gene-related peptide (CGRP), have long been described to exert pro- and anti-inflammatory effects respectively (6–8). For example, SP is well established to increase blood vessel dilation and permeability, in addition to increasing adhesion molecule expression on endothelial cells (7, 9, 10). These physiological changes that allow for increased immune cell recruitment into the local tissue environment are the basis of neurogenic inflammation. These immunologically potent neuropeptides from sensory neurons have a critical role in the response to infection with bacterial pathogens of the lung, skin, small intestine, and colon.

Our previous studies demonstrated that the host response to the mouse-adapted attaching and effacing (A/E) bacterial pathogen *Citrobacter rodentium* was significantly reduced in mice with prior ablation of TRPV1^+^ sensory neurons (11). This enteric bacterial pathogen possesses virulence genes encoded by the locus of attachment and effacement in a pathogenicity island, a type 3 secretion system, and injects bacterial effector proteins (12, 13). These key aspects have made this model fundamental to our understanding of the host cellular and molecular responses to A/E pathogens *in vivo*. Infection with *C. rodentium* triggers host-protective IL-22 production from innate lymphoid cells with the recruitment of monocytes and neutrophils early in the course of infection until adaptive immunity responses are generated (14–17). Adaptive immune responses during *C. rodentium* infection are characterized by the recruitment of CD4^+^ T cells recruitment that produce IFNγ, IL-17A, and IL-22 and B cells which are required for clearance of the pathogen (18–20). Previously, we demonstrated that TRPV1^+^ neurons aided the coordination of this host response (11); however, the precise role of TRPV1 in the host response during *C. rodentium* infection was not assessed. Purported expression of TRPV1 has been reported on numerous cell types in the intestinal tract in addition to neurons ranging from intestinal epithelial cells to T cells (21, 22). Perhaps it is unsurprising that both pro- and anti-inflammatory roles for TRPV1 have been described in models of intestinal inflammation (23). It is uncertain if these seemingly contradictory data are simply reflective of the biological complexity or the unique models of intestinal inflammation. Here we assessed the role of TRPV1 in the development of host immune responses during *C. rodentium* infection, using wildtype (WT) and TRPV1 knockout (TRPV1^-/-^) mice.

Deficiency in TRPV1 significantly increased the bacterial burden at the peak of infection. The increased *C. rodentium* burden in TRPV1^-/-^ mice caused increased colonic inflammation and colonic crypt hyperplasia, which was not due to reduced expression of IFNγ, IL-17A, or IL-22, or the recruitment of T cells into the colon producing these host protective cytokines compared to WT infected mice. Interestingly, infection of TRPV1^-/-^ mice with C. rodentium resulted in significantly fewer colonic neutrophils 10 days post-infection compared to WT mice. This was corroborated by a reduction in expression of the neutrophil chemokine *Cxcl6*. Reduced recruitment was not due to a defect in neutrophil maturation in the bone marrow, but instead driven by downregulation of cell adhesion molecules necessary for rolling adhesion and extravasation. Flow cytometry analysis of colonic blood endothelial cells revealed a significant decrease in ICAM-1 and VCAM-1, but not MAdCAM-1 expression in TRPV1^-/-^ mice compared to WT mice. Collectively these data identify a novel role for TRPV1 in the regulation of neutrophil recruitment to the colon during infection with enteric bacterial pathogens.

## Methods

### Animals

C57BL/6 (wildtype, WT) and TRPV1^-/-^ mice were originally purchased from The Jackson Laboratory (Bar Harbor, ME) to establish a breeding colony in our vivarium. Male and female TRPV1^-/-^ mice and C57BL/6 mice were maintained in a specific pathogen-free environment and used for experiments at 6 to 8 weeks old. All animals had *ad libitum* access to food and water. All procedures were approved by the institutional animal care and use committee at UC Davis, in accordance with the Guide for Care and Use of Laboratory Animals. Mice were euthanized by CO_2_ asphyxiation followed by cervical dislocation according to American Veterinary Medical Association guidelines.

### *Citrobacter rodentium* and bacterial burden quantification

*Citrobacter rodentium*, strain DBS100, was generously provided by Dr. Andreas Baumler (UC Davis, Davis, CA). Bacteria were grown from frozen stocks on MacConkey agar at 37°C and a single colony was grown in LB broth overnight at 37°C. The bacterial suspension was re-grown at 37°C to reach a final infection suspension containing 10^8^ colony-forming units (CFU). Mice were inoculated by oral gavage with 0.1 mL of the bacterial suspension or sterile LB broth. Mice were euthanized on day 10 or 29 post-infection for tissue analyses. *C. rodentium* colonization was quantified on day 10, 29 by homogenization of either fecal pellet or distal colonic tissue in 1 mL PBS and plating serial dilutions onto MacConkey agar. *C. rodentium* colonies were counted after overnight growth at 37°C and results are expressed as CFU/g feces or colonic tissue.

### Histology

Distal colon (1 cm) sections were fixed in 10% buffered formalin, and paraffin embedded for cross-sectioning. Sections (6 µm) were cut and stained with hematoxylin and eosin (H&E). Epithelial cell hyperplasia was evaluated using bright-field microscopy at 20X objective, by measuring the crypt length of 20 well-oriented colonic crypts for each mouse using FIJI (Fiji is just ImageJ, NIH).

### Quantitative PCR

Gene expression was measured by quantitative real-time PCR as previously described (11). Briefly, tissues were homogenized in TRIzol using a bead beater allowing for isolation of RNA. This RNA was then used to prepare cDNA by reverse transcription (iScript, Bio-Rad Hercules CA) in order to conduct real-time PCR using the indicated primer pairs from PrimerBank (24) **(Table S1)** with SYBR Green master mix (ThermoFisher, Waltham MA).

### Immunohistochemistry and confocal microscopy

Paraffin sections of colonic tissue (6 µm) were used for confocal analysis with antibodies raised against specific proteins of interest according to standard protocols (25). In brief, after slides were de-paraffinized and rehydrated, antigen retrieval was performed in citrate buffer (10 mM, pH 6.0, 30 min., 95°C). After blocking in 5% BSA (w/v) and normal goat serum, samples were incubated in primary antibody overnight (16 h, 4°C). Slides were washed extensively (3 x 5 mins) in TBS-tween20 and incubated in appropriately labeled secondary antibodies (Invitrogen) for 1 h at room temperature, washed, counterstained with DAPI in TBS-tritonX100 0.1% v/v, washed and mounted in Prolong gold (ThermoFisher, Waltham, MA). Staining using anti-mouse CDH1 (E-cadherin) and anti-mouse βIII Tubulin **(Table S2)** was revealed using a mouse-on-mouse kit according to manufacturer’s instructions (Vector laboratories, Burlingame, CA).

Primary and secondary antibodies used are detailed in **Table S2.** Slides were imaged on a Leica SP8 STED 3X confocal microscope with a 40X 1.3NA objective or a 63X 1.4 NA objective. All areas larger than the field of view of the objective were acquired using a tiling approach, whereby adjacent images were acquired with a 10% overlap, and processed by Imaris Stitcher (Oxford Instruments United Kingdom).

### T Cell Enrichment

Inguinal and mesenteric lymph nodes and spleen were sterilely excised from non-treated C57BL/6 mice and placed on a 100μm filter and dissociated using the plunger of a syringe followed by several washing steps with stain buffer (1X PBS + 2% FBS). Single cell suspension was treated with Ack lysis solution for 5 minutes at room temperature before being washed.

Cells were then incubated with Fc block (anti-CD16/32, 10 µg/ml, Tonbo Biosciences, San Diego, CA) for 15-minute on ice. Antibody cocktail was added to surface stain cells for 30 minutes on ice, followed by washing in stain buffer. The following biotinylated antibodies were added to this cocktail at 1:50 dilution: anti-CD161 (clone# PK136, Ref# 30-5941), anti-CD11c (clone# N418, Ref# 30-0114), anti-Ly6G (Clone# RB6-8C5, Ref# 30-5931), anti-TER119 (Clone# TER-119, Ref# 30-5921), anti-CD11b (Clone# M1/70, Ref# 30-0112), and anti-B220 (Clone# RA3-6B2, Ref# 30-0452) from Tonbo Biosciences. Cells were then washed and resuspended in magnetic streptavidin beads (Cat# 557812, BD Biosciences, Franklin Lakes NJ according to the manufacturer’s protocol. Cells were incubated under mild rotation at 4°C for 30 minutes and then the solution volume was brought up to 1mL total. Tube was placed in BD IMAG Cell Separation Magnet (Cat# 552311 BD Biosciences, Franklin Lakes NJ) for 8 minutes and the negative fraction was taken and placed into a fresh tube on the IMAG. This was again incubated for 6 minutes and repeated one additional time. All negative fractions were combined, and a fraction of the cells were stained for flow cytometry analysis to confirm T cell purity was above 90%.

### T Cell Proliferation Assay

96-well round bottom plates were coated with (0, 1, 10 μg/mL) anti-CD3ε (clone# 17A2, Ref# 40-0032, Tonbo Biosciences, San Diego, CA) or PBS for 2 hours at 37°C and removed by aspiration. IMAG negatively selected CD4^+^ T Cells were dyed with CellProliferation Dye eFluor450 (Ref# 65-0842-85, eBioscience, San Diego, CA) according to manufacturer’s protocol then plated in 96-well round bottom plates and cultured in the presence of anti-CD28 (0, 4, 10 μg/mL, Ref# 40-0281, Tonbo Biosciences, San Diego, CA) and/ or capsaicin. Cells were incubated in RPMI media with 10% FBS, 2mM L-glutamine, 50 µM β-mercaptoethanol, and 1% Penicillin/ Streptomycin statically at 37°C 5% CO_2_ for 72 hours before being harvested for flow cytometry analysis. Proliferation index was determined using the Cell Proliferation Cycle tool in FlowJo.

### Isolation of cells from the colon

Lamina propria lymphocytes were isolated using a lamina propria dissociation kit in combination with a gentleMACS tissue dissociator according to the manufacturer’s instructions (Miltenyi Biotec, Gaithersburg, MD). In brief, colons were removed from the euthanized mouse, opened longitudinally, and cut into 0.5 cm long segments. Epithelial cells were removed by gentle agitation (200 RPM) of tissue fragments in HBSS supplemented with 5 mM EDTA and 5% FBS solution (without Ca^2+^ and Mg^2+^), for 20 minutes at 37°C. Tissue fragments were transferred to a gentleMACS C-tube, with HBSS (with Ca^2+^ and Mg^2+^) and MACS digestion enzymes, and incubated while shaking for 30 minutes at 37°C, followed by dissociation by the gentleMACS device. The resulting single cell suspension was passed through a 100 µm strainer, washed extensively, and subjected to staining.

### Bone Marrow Cell Isolation

Wild-type and TRPV1^-/-^ euthanized mice had right femur removed. Bone was isolated by removing skin, muscle, and fur and then rinsed with PBS. Ends of the bone were cut and a 26-gauge needle and syringe were used to push PBS through the bone and remove the marrow. Cells were then treated with Ack lysis buffer for 5 minutes at room temperature before undergoing cell surface staining.

### Flow cytometry

Staining of cells was performed using a standard protocol. In brief, cells were counted manually by hemocytometer with trypan blue exclusion and dispensed into flow cytometry tubes, centrifuged, and resuspended in staining buffer containing Fc block (anti-CD16/32, 10 µg/ml, Tonbo Biosciences, San Diego, CA) for 25-minute on ice. Antibody cocktail **(Table S3)** was added to surface stain cells for 30 minutes on ice, followed by washing in stain buffer. Viability was determined using live/dead aqua according to manufacturer’s instructions (ThermoFisher, Waltham MA). All flow cytometry data was acquired on a LSRII (BD Biosciences, Franklin Lakes NJ) using DIVA software, with analysis using FlowJo (Treestar, Eugene OR)

### Intracellular staining

Prior to surface staining, cells were incubated in RPMI media with 10% FBS, 1% Penicillin/ Streptomycin, 2mM L-glutamine, and BD GolgiPlug (1:500, Cat# 555029, BD Biosciences, Franklin Lakes NJ) for 4 hours at 37°C. To stimulate cells, 1X Cell Stimulation Cocktail (phorbol 12-myristate 13-acetate) was added (eBioscience, San Diego CA). Following surface staining cells were fixed and permeabilized using a BD Cytofix/ Cytoperm Fixation/ Permeabilization Solution kit (Cat# 554714 BD Biosciences, Franklin Lakes NJ) followed by intracellular staining with anti-IFNγ, anti-IL-17a, and anti-IL-22 **(Table S3)** in 1X BD Perm/ Wash Buffer (Cat# 554723 BD Biosciences, Franklin Lakes NJ).

### Ussing Chamber

Mouse colon was excised, cut along the mesenteric border, and then mounted onto cassettes (Physiologic Instruments) allowing for 0.1 cm^2^ of the mounted colonic tissue to be exposed to 4 mL of circulating oxygenated Ringer’s buffer (115 mM NaCl, 1.25 mM CaCl_2_, 1.2 mM MgCl_2_, 2.0 mM KH_2_PO_4_, and 25 mM NaHCO_3_) at 37°C. To keep the colonic tissues healthy and viable throughout the experiment, 10 mM glucose (Sigma Aldrich, St. Louis, MO) was added to both the serosal and mucosal compartments. Two pairs of electrodes attached to agar-salt bridges were used to monitor the potential difference between both compartments and to inject current during voltage clamping. Recordings and current injection were performed using an automated voltage clamp and Acquire and Analyze software (Physiologic Instruments, San Diego CA).

After 20 minutes of equilibration, 88 mg/mL of FITC-labeled dextran (Sigma Aldrich, St. Louis, MO) was added to the mucosal chamber to measure tight junction permeability over time. This was achieved by collecting samples from the serosal compartments every 30 minutes for 2 hours and measuring the respective FITC concentrations on a plate reader (Synergy H1; BioTek, Winooski, VT). Samples with visible FITC leakage in the serosal compartment were omitted. Baseline active ion transport was measured by short-circuit current (Isc) and tight junction permeability was measured by conductance (G).

### Epithelial Organoid Cell Culture & Proliferation

Colonic crypts were isolated and cultured as described previously with minor modifications (26). Mouse colon was dissected out, flushed thoroughly with ice cold PBS, opened longitudinally and then cut into 5 mm-long fragments. Tissue fragments were washed repeatedly in ice-cold PBS and then incubated in chelation buffer (5.6 mM Na_2_HPO_4_, 8 mM KH_2_PO_4_, 96 mM NaCl, 16 mM KCl, 44 mM sucrose, 54.8 mM D-sorbitol, 5 mM EDTA and 0.5 mM DTT) for 30 minutes on ice. Colonic crypts were gradually released from the tissue fragments during 12 rounds of washing in 2% fetal bovine serum and filtering through 70 μm filter mesh, generating 12 fractions. Only fractions 8 to 12 containing enriched colonic crypts were used for downstream experiments.

About 200 colonic crypts were seeded in 25 μL Cultrex BME Type 2 (R&D systems, Minneapolis, MN) domes and cultured in Advanced DMEM/ F-12 supplemented with N2, B27, 50 ng/mL EGF, 100 ng/mL Noggin, 500 ng/mL R-spondin, 100 ng/mL Wnt-3an, 1 mM N-acetyl-L-cysteine and 1 μM ROCK inhibitor (Y27632, Millipore, Burlington, MA). Organoid proliferation was assessed with EdU incorporation assay at day 5 post crypt seeding. EdU (10 μΜ) was added to the culture media for 24 h and visualization was performed according to the manufacturer’s instruction with Click-iT EdU Cell Proliferation Kit Alexa Fluor 647 (ThermoFisher, Waltham, MA). Organoids were imaged on a Leica SP8 STED 3X confocal microscope with a 40X objective. The percentage of EdU positive cells was quantified manually with Imaris software.

### Statistics

Statistical analysis of all data was performed using Prism 9.0 (GraphPad, La Jolla, CA) with a Student’s t test or one-way or two-way ANOVA followed by a post-hoc analysis with Tukey’s multiple comparison test. Individual data points are presented as mean ± standard error of the mean.

## Results

### TRPV1 deficiency increases bacterial burden and colonic pathology during *C. rodentium* infection

To determine the role of TRPV1 in the host response to an A/E pathogen *in vivo*, TRPV1^-/-^ and WT mice were administered LB (control) or C. *rodentium* by orogastric gavage. Fecal and colonic *C. rodentium* were significantly increased in TRPV1^-/-^ compared to WT mice 10 days post-infection (p.i.) (**Figure 1A**), with both TRPV1^-/-^ and WT mice clearing the infection by 29 days p.i. (**Figure 1B**). This significant increase in bacterial burden 10 days p.i. in TRPV1^-/-^ mice was associated with increased severity of *C. rodentium* induced colonic pathology. As expected, colonic crypt hyperplasia was observed in mice infected with *C. rodentium* compared to uninfected controls. This increase was significantly exacerbated in TRPV1^-/-^ compared to WT mice at 10- and 29-days p.i. (**Figure 1C & 1D)**. Infection-induced crypt hyperplasia was due to increased intestinal epithelial cell proliferation as revealed by significantly increased Ki67^+^ CDH1^+^ DAPI^+^ cells 10- and 29-days p.i. (**Figure 1E & 1F)**. Small but significantly increased crypt length were observed in uninfected TRPV1^-/-^ compared to WT mice, though there was no increased basal intestinal epithelial cell (IEC) proliferation in uninfected mice.

**Fig. 1.**
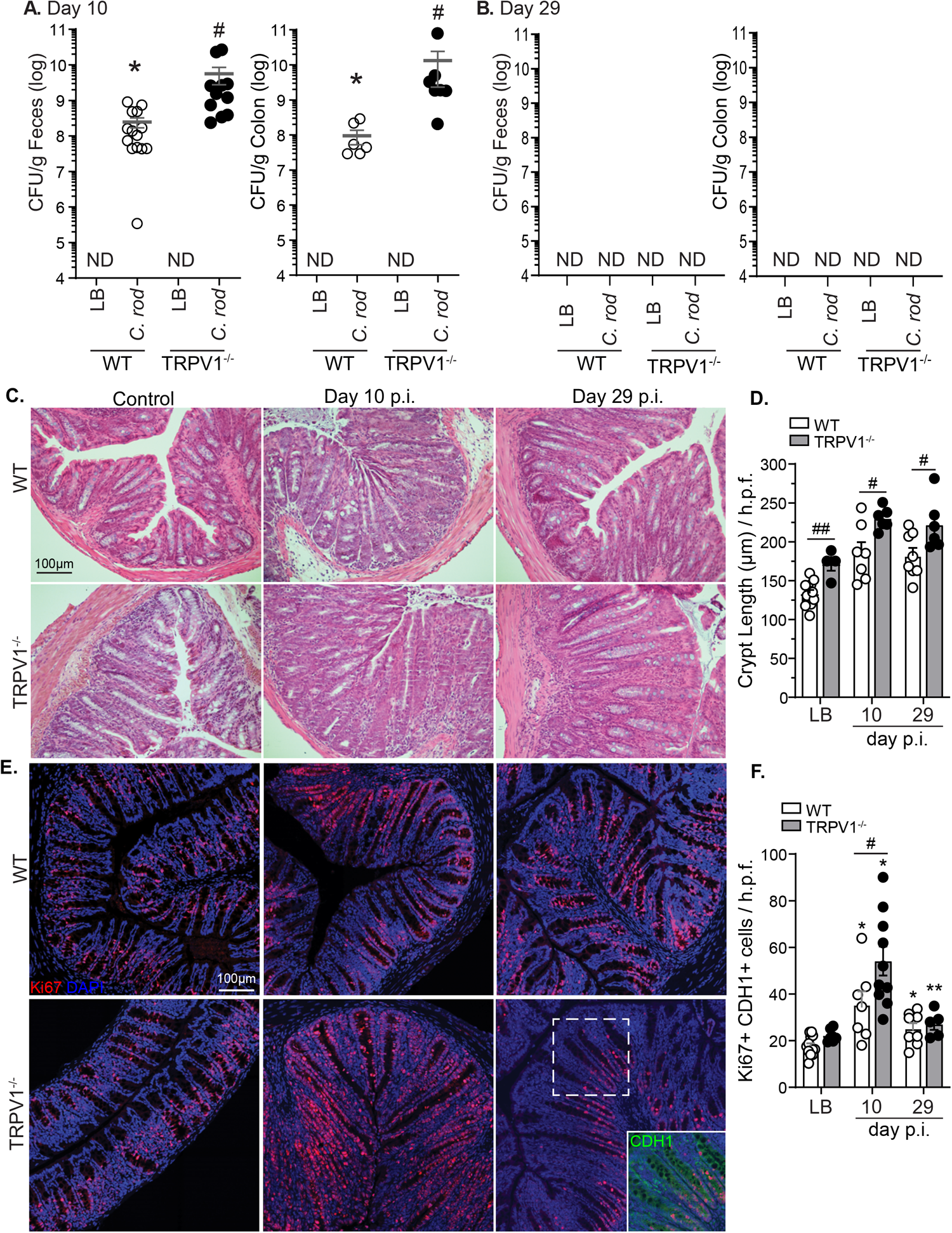
TRPV1^-/-^ mice have an increased bacterial burden and colonic crypt hyperplasia during *C. rodentium* infection. Wild-type (WT; open circles) and TRPV1^-/-^ mice (black circles) were infected with *C. rodentium* or given vehicle control (LB) by oral gavage. The number of fecal and colonic tissue adherent bacteria were assessed in WT and TRPV1^-/-^ mice at **(A)** 10 days p.i. and **(B)** 29 days p.i. **(C)** Hematoxylin and eosin (H&E) stained paraffin-embedded cross-sections of colon tissue from non-infected control, or 10, 29 days p.i. (Scale bar: 100µm), with **(D)** crypt length measured using FIJI. **(E)** Confocal images of distal colon tissue from non-infected, 10, or 29 days p.i. mice stained with Ki67 (red), CDH1 (green), and DAPI (blue), with **(F)** quantification of Ki67^+^ CDH1^+^ DAPI^+^ cells. Data are from 7-12 mice per group in 4 separate experiments, and presented as mean ± standard error of the mean: *, *P* < 0.05, **, *P* < 0.01, and ***, *P* < 0.001 vs uninfected mice; #, *P* < .05, # #, P< 0.01 compared with WT mice; Student’s *t* test **(A & B)** and two-way ANOVA **(D & F)** with post-hoc analysis using Tukey’s multiple comparisons test. LB, Luria-Bertai; p.i., post-infection.

Although expression of TRPV1 has been reported on many cell types including sensory neurons, intestinal epithelial cells, and immune cells (27–31), immunostaining revealed that TRPV1 immunoreactivity was retained in TRPV1-/- mice and was associated with epithelial cells. In fact, neuronal TRPV1 immunoreactivity (βIII-tubulin^+^) was only found in WT and not TRPV1^-/-^ colon **(Figure S1A)**. With potential for TRPV1 expression on intestinal epithelial cells, we investigated the impact of TRPV1 on epithelial cell proliferation and function. Assessment of colonic permeability by FITC-dextran in Ussing chambers found no significant increase in permeability of TRPV1^-/-^ vs WT mice **(Figure S1B)**. Intestinal epithelial cell proliferation was further found not to be dependent on TRPV1, as incorporation of the thymidine analogue EdU and organoid diameter was found to be equivalent in WT and TRPV1^-/-^ organoids at varied concentrations of the TRPV1 agonist, capsaicin **(Figure S1C)**.

### CD4+ T cell response to *C. rodentium* is intact in TRPV1^-/-^ mice

It is well established that the differentiation and recruitment of specific CD4^+^ T cell subsets are required for the control of *C. rodentium* infection (32, 33). As TRPV1 has been suggested to enhance proliferation and differentiation of T cells in a cell intrinsic manner (34), we assessed if T cell responses during *C. rodentium* infection were altered in TRPV1^-/-^ compared to WT mice. Enumeration of colonic T cells revealed no significant differences in TRPV1^-/-^ vs WT mice by confocal microscopy after 10- and 29-days p.i. (**Figure 2A & 2B)**. These data were confirmed by flow cytometry analyzing the number of CD3^+^ CD4^+^ T cells present in the lamina propria at baseline and 10-days p.i. (**Figure 2C**). Intracellular cytokine staining conducted on these lamina propria lymphocytes further revealed no significant difference in the frequency of CD3^+^ CD4^+^ T cells producing IFNγ and IL-22, with a slight but significant decrease in the frequency of IL-17A+ T cells in TRPV1^-/-^ compared to WT mice 10 days p.i. (**Figure 2D-F**). Quantification by qPCR revealed no significant difference in *IFNγ*, *IL-17a*, or *IL-22* expression in the colon of WT or TRPV1^-/-^ mice 10- or 29-days p.i. **(Figure S2A-C)**. To investigate if TRPV1 could act in a cell intrinsic manner to regulate T cell proliferation, we assessed the proliferative response of WT and TRPV1^-/-^ CD4^+^ T cells. Stimulation with increasing concentrations of anti-CD3ε, anti-CD28, with and without the TRPV1 agonist capsaicin showed no difference in proliferation **(Figure S2D & S2E)**. Together, these data demonstrate that T cell responses in TRPV1^-/-^ mice remain intact during *C. rodentium* infection.

**Fig. 2.**
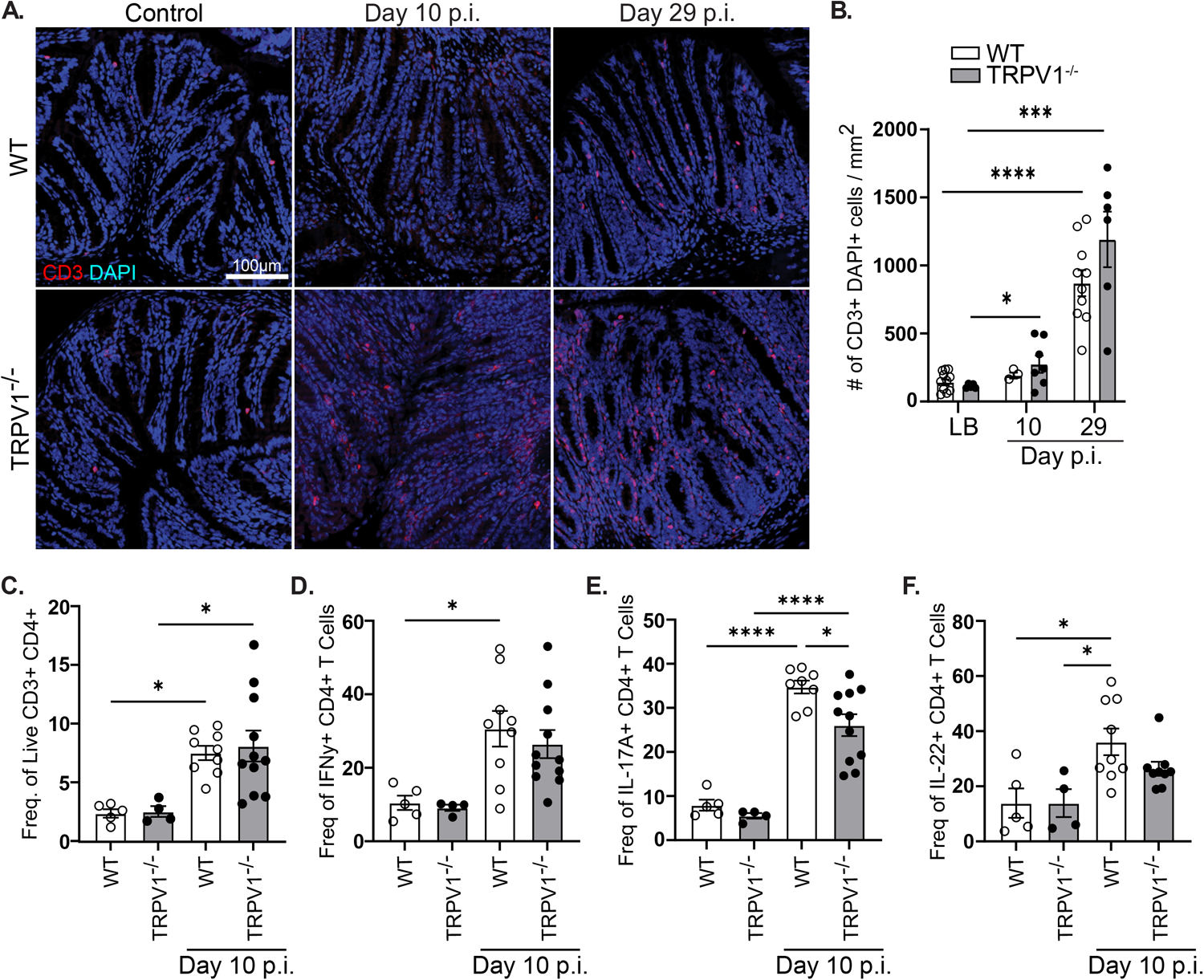
T cell recruitment and cytokine production are unaffected by TRPV1 deficiency during *C. rodentium* infection. **(A & B)** Colonic tissue sections were assessed for CD3^+^ (red) DAPI^+^ (blue) T cell infiltration in vehicle control (LB) and *C. rodentium* infected wild-type (WT) and TRPV1^-/-^ mice after 10- or 29-days p.i. **(C)** Lamina propria lymphocytes were assessed by flow cytometry to enumerate live CD45^+^ CD3^+^ CD4^+^ T cells and determine the frequency of **(D)** IFNγ^+^, **(E)** IL-17A^+^, or **F)** IL-22^+^ T cells in control and infected WT and TRPV1^-/-^ mice 10 days p.i. Data are presented as mean ± standard error of the mean: *, P < 0.05, **, P < 0.01 and ***, P < 0.001; one-way ANOVA with post-hoc analysis using Tukey’s multiple comparisons test, with 4– 11 animals per group. LB, Luria-Bertai; p.i., post-infection.

### Lack of TRPV1 causes dysregulation of innate immune responses during *C. rodentium* infection

With the increased bacterial burden and histopathology in *C. rodentium* infected TRPV1^-/-^ mice compared to WT mice without reduced T cell recruitment or cytokine expression, we sought to characterize the host innate immune response. Infection of WT and TRPV1^-/-^ mice significantly increased colonic *Il6* and *Tnfα* expression compared to uninfected controls (**Figure 3A & 3B)**. While *Il6* mRNA levels returned to baseline by day 29 p.i. in WT, significantly increased *Il6* was still observed in TRPV1^-/-^ mice at this time (**Figure 3A**). Expression of *Il1β* was significantly decreased in TRPV1^-/-^ mice at 10 days p.i. compared to WT, but both WT and TRPV1^-/-^ mice returned to basal expression by 29 days p.i. (**Figure 3C**).

**Fig. 3.**
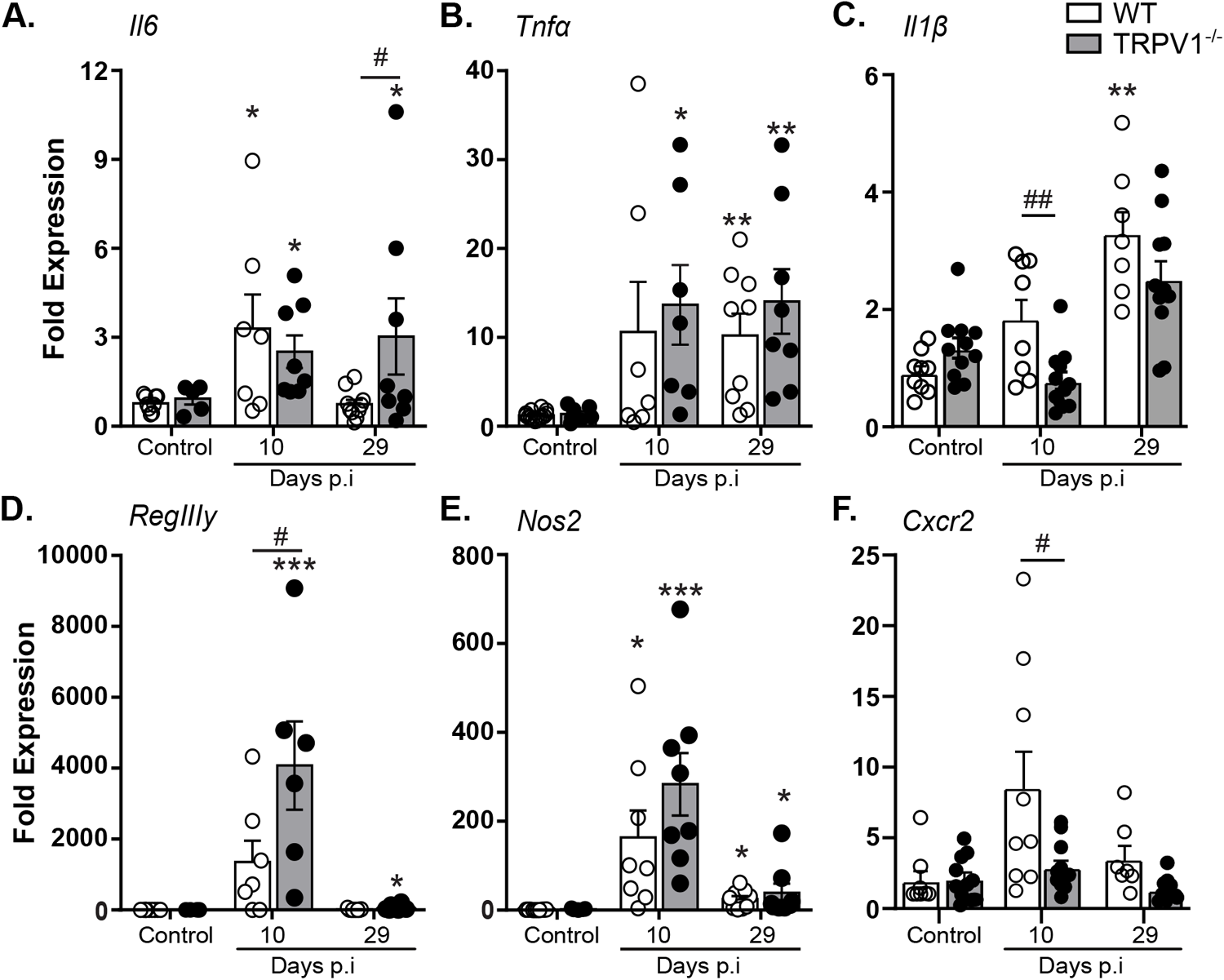
Select innate immune responses are reduced in *C. rodentium*-infected TRPV1^-/-^ mice. The host immune response during infection was assessed through qPCR conducted on colonic tissue from control or *C. rodentium* infected wild-type (WT) or TRPV1^-/-^ mice. These include the expression of proinflammatory cytokines **(A)** *Il6*, **(B)** *Tnfα*, **(C)** *Il1β,* **(D)** the antimicrobial peptide *RegIIIy*, **(E)** inducible nitric oxide synthase (*Nos2*), and **(F)** the neutrophil chemokine receptor *Cxcr2.* Data are presented as mean ± standard error of the mean, n= 7-13 animals/ group: *, P < 0.05, **, P < 0.01, and ***, P < 0.0001 versus uninfected mice; #, P < 0.05 and ###, P < 0.0001 compared with WT *C. rodentium*-infected mice; one-way ANOVA with post-hoc analysis using Tukey’s multiple comparisons test. p.i., post-infection.

Reflective of the increased *C. rodentium* burden, expression of the antimicrobial gene RegIIIγ was significantly increased during infection, and further enhanced in infected TRPV1^-/-^ compared to WT mice 10 days p.i. (**Figure 3D**). However, altered expression of innate and host protective genes in *C. rodentium* infected TRPV1^-/-^ mice was not universal. Although inducible nitric oxide synthase (iNOS, NOS2) expression was induced by *C. rodentium* infection, no difference was detected 10- or 29-days p.i. in WT and TRPV1^-/-^ mice (**Figure 3E**). Interestingly, we found a significant decrease in the expression of *Cxcr2*, a neutrophil chemokine receptor critical for chemotaxis into the lumen of the gut, in TRPV1^-/-^ mice compared to WT at 10 days p.i. (**Figure 3F**). These data suggest neutrophil recruitment may be altered in TRPV1^-/-^ mice during *C. rodentium* infection.

### Colonic neutrophil recruitment during *C. rodentium* infection is reduced in TRPV1^-/-^ mice

To determine the effect of TRPV1 on neutrophil recruitment to the colon during *C. rodentium* infection, we first assessed colonic expression of chemokines in TRPV1^-/-^ and WT mice. Expression of *Cxcl1* was significantly increased at 10 days p.i. irrespective of genotype. Similarly, *Cxcl2 and Cxcl3* expression were increased during infection; however, there was a significant decrease of these chemokines 29 days p.i. in TPRV1^-/-^ compared to WT mice. *Cxcl6*, a critical neutrophil chemokine and antimicrobial peptide (35), was significantly decreased in TRPV1^-/-^ compared to WT mice at 10- and 29-days p.i. (**Figure 4A**). Given these decreases in chemokines that recruit neutrophils, and reduced colonic *Cxcr2*, we assessed neutrophil recruitment to the colonic lamina propria during *C. rodentium* infection in TRPV1^-/-^ and WT mice. Flow cytometry on colonic lamina propria demonstrated a significant decrease in number of and frequency of live neutrophils in the colon at 10 days p.i. in TRPV1^-/-^ mice compared to WT (**Figure 4B-D**); however, other innate immune cell populations such as monocytes, macrophages, or conventional dendritic cells were not impacted (**Figure 4E-G**). Despite differences in *Cxcl6*, neutrophil chemokines are often redundant and conserved, so loss of one may not impact neutrophil chemotaxis entirely on its own (36, 37). To determine if the significant reductions in colonic neutrophils were due to reduced differentiation, pre-neutrophil, immature neutrophil, and mature neutrophil subsets within the bone marrow at baseline and 10 days p.i. were assessed and found no significant difference between WT and TRPV1^-/-^ mice **(Figure S4B)**. Together, these data demonstrate that the TRPV1 induced reduction of colonic neutrophil recruitment during *C. rodentium* infection is not due to cell maturation.

**Fig. 4.**
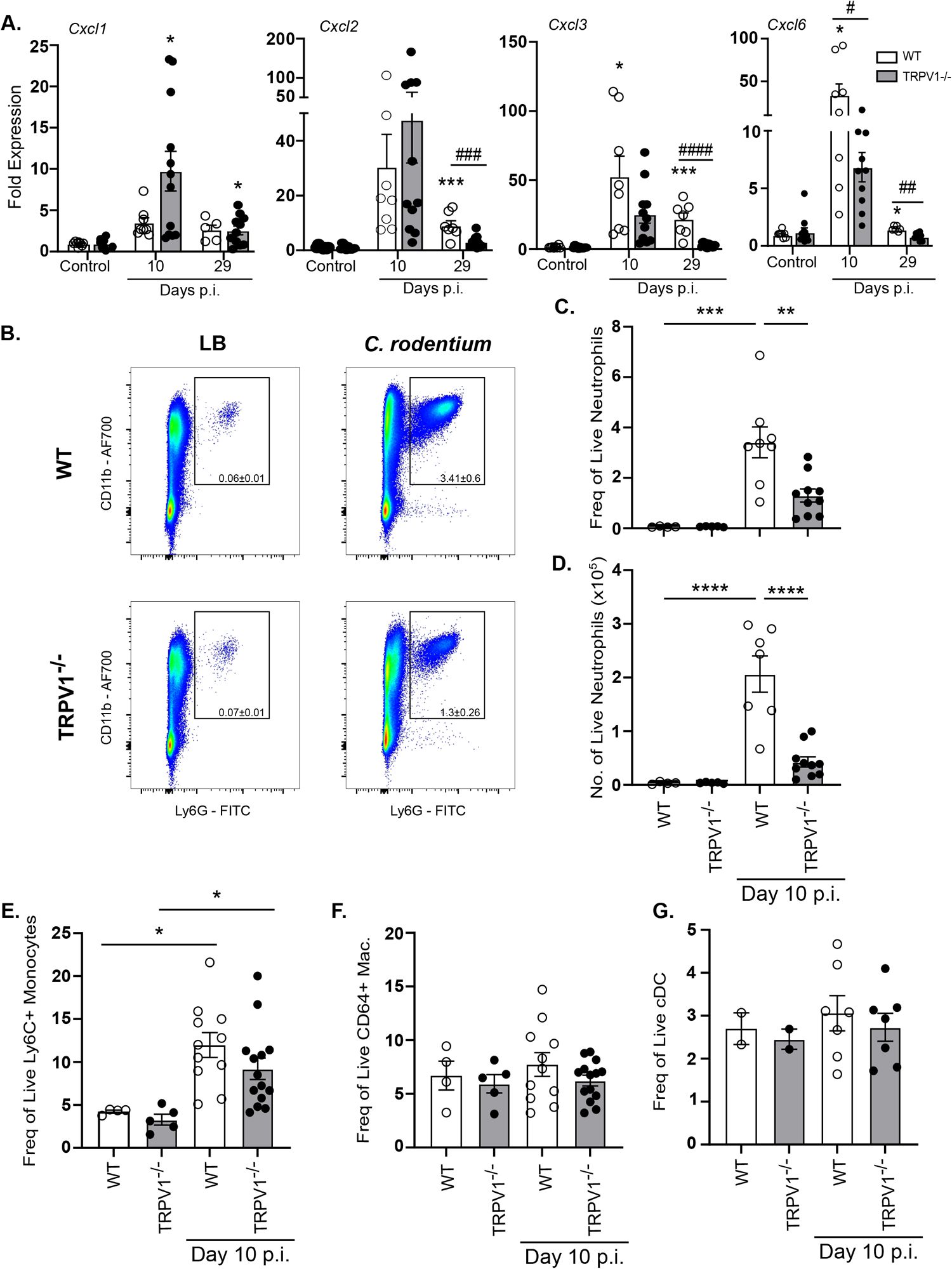
Neutrophil recruitment is impaired in TRPV1^-/-^ mice during *C. rodentium* infection. **(A)** Common neutrophil chemokines *Cxcl1*, *Cxcl2*, *Cxcl3*, and *Cxcl6* were assessed by qPCR performed on colonic tissue from control or *C. rodentium* infected wild-type (WT) or TRPV1^-/-^ mice. **(B-D)** Neutrophil recruitment was determined by flow cytometry for live CD45^+^ CD11b^+^ Ly6G^+^ cells in the whole colon after epithelial cells were removed from WT and TRPV1^-/-^ mice given vehicle control (LB) and 10 days p.i. with *C. rodentium* **(B)** Representative flow plots and cumulative data of **(C)** frequency of live neutrophils and **(D)** total cell number of live neutrophils in the colon. **(E-G)** Cumulative data of frequency of live **(E)** Ly6C^+^ monocytes, **(F)** CD64^+^ macrophages, and **(G)** CD11c^hi^ conventional dendritic cells (cDC) in the lamina propria of the colon of WT and TRPV1^-/-^ mice at baseline and 10 days p.i. Data are presented as mean ± standard error of the mean: *, P < 0.05, **, P < 0.01 and ***, P < 0.001 compared with uninfected mice; #, P < 0.05 compared with WT *C. rodentium*-infected mice **(A)** *, P < 0.05, **, P < 0.01, and ***, P < 0.0001 **(C-G)**; one-way ANOVA with post-hoc analysis using Tukey’s multiple comparisons test. 4–11 animals per group. LB, Luria-Bertani; p.i., post-infection.

### TRPV1 regulates colonic blood endothelial cell expression of selective adhesion molecules during *C. rodentium* infection

Considering the reduced numbers of colonic neutrophils that were not due to development of neutrophils in bone marrow, we assessed if TRPV1^-/-^ mice had deficiencies in the processes that regulated neutrophil recruitment. Using flow cytometry, *C. rodentium* infection (10 days p.i.) was found to significantly increase ICAM-1 and VCAM-1 on the surface of colonic lamina propria blood endothelial cells (single, live, CD45^-^, CD31^+^, gp38^-^) in WT but critically not in TRPV1^-/-^ mice. Infection induced changes in adhesion molecule expression were not global, as MAdCAM-1^+^ blood endothelial cells were not different in WT or TRPV1^-/-^ mice. (**Figure 5A &B)**. Quantification of mRNA transcripts by qPCR for these cell adhesion molecules revealed that *Icam1* but not *Vcam1* or *Madcam1* was significantly decreased at 10 days p.i. in TRPV1^-/-^ mice compared to WT (**Figure 5C**). Together these data suggest that TRPV1 regulates the recruitment of immune cells through regulation of adhesion molecules expressed by blood endothelial cells in the colon.

**Fig. 5.**
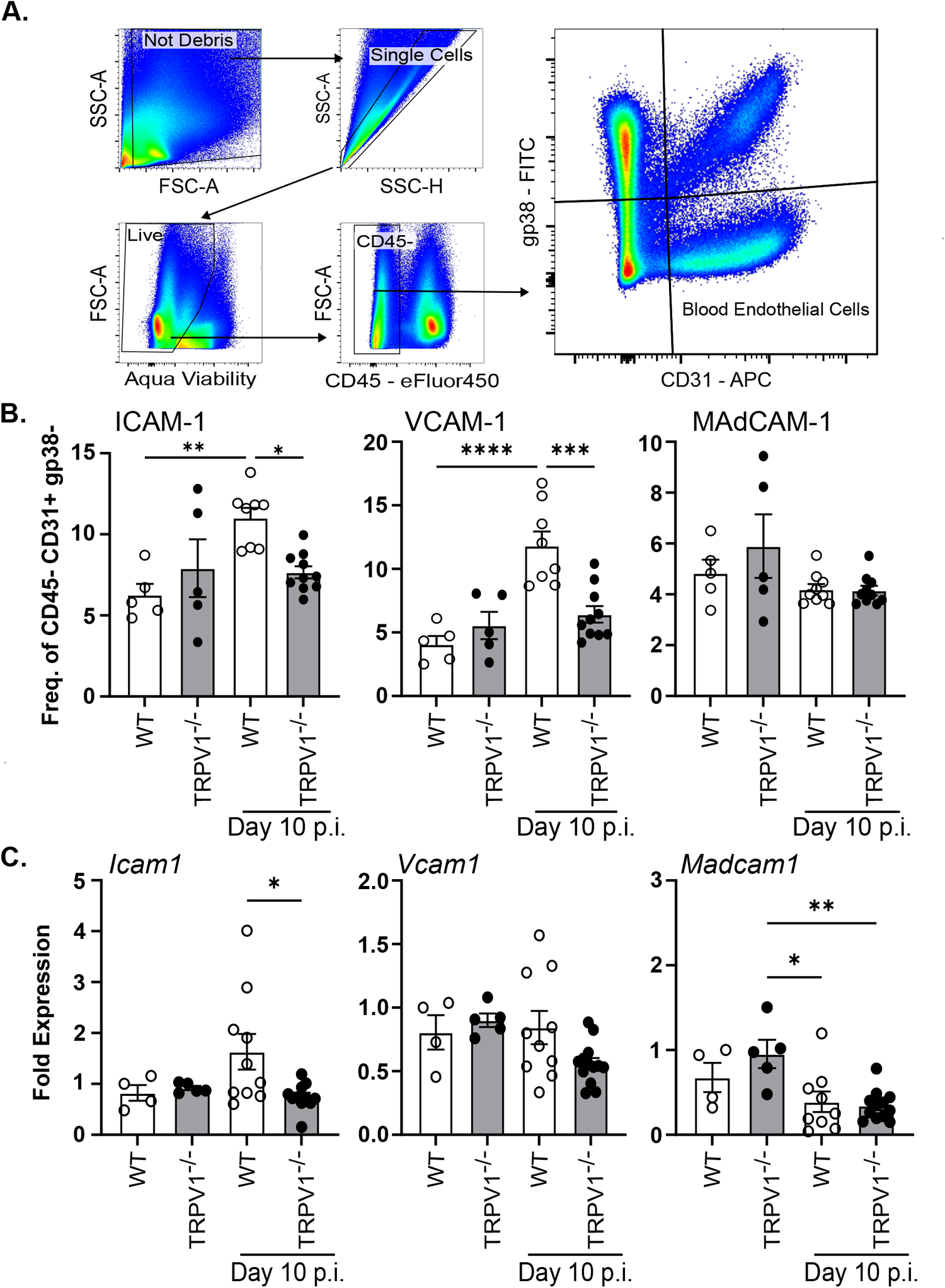
TRPV1 dependent neutrophil recruitment is driven by upregulation of ICAM-1 and VCAM-1 on blood endothelial cells. **(A & B)** Colonic blood endothelial cells from wild-type (WT) and TRPV1^-/-^ mice were assessed for their expression of ICAM-1, VCAM-1, and MAdCAM-1 by flow cytometry at baseline and 10 days p.i. (C) *Icam1*, *Vcam1*, and *Madcam1* mRNA were assessed by qPCR of the colonic tissue at baseline and 10 days p.i. Data are presented as mean ± standard error of the mean: *, P < 0.05, **, P < 0.01 and ***, P < 0.001; one-way ANOVA with post-hoc analysis using Tukey’s multiple comparisons test. 4–10 animals per group. LB, Luria-Bertani; p.i., post-infection.

**Fig. 6.**
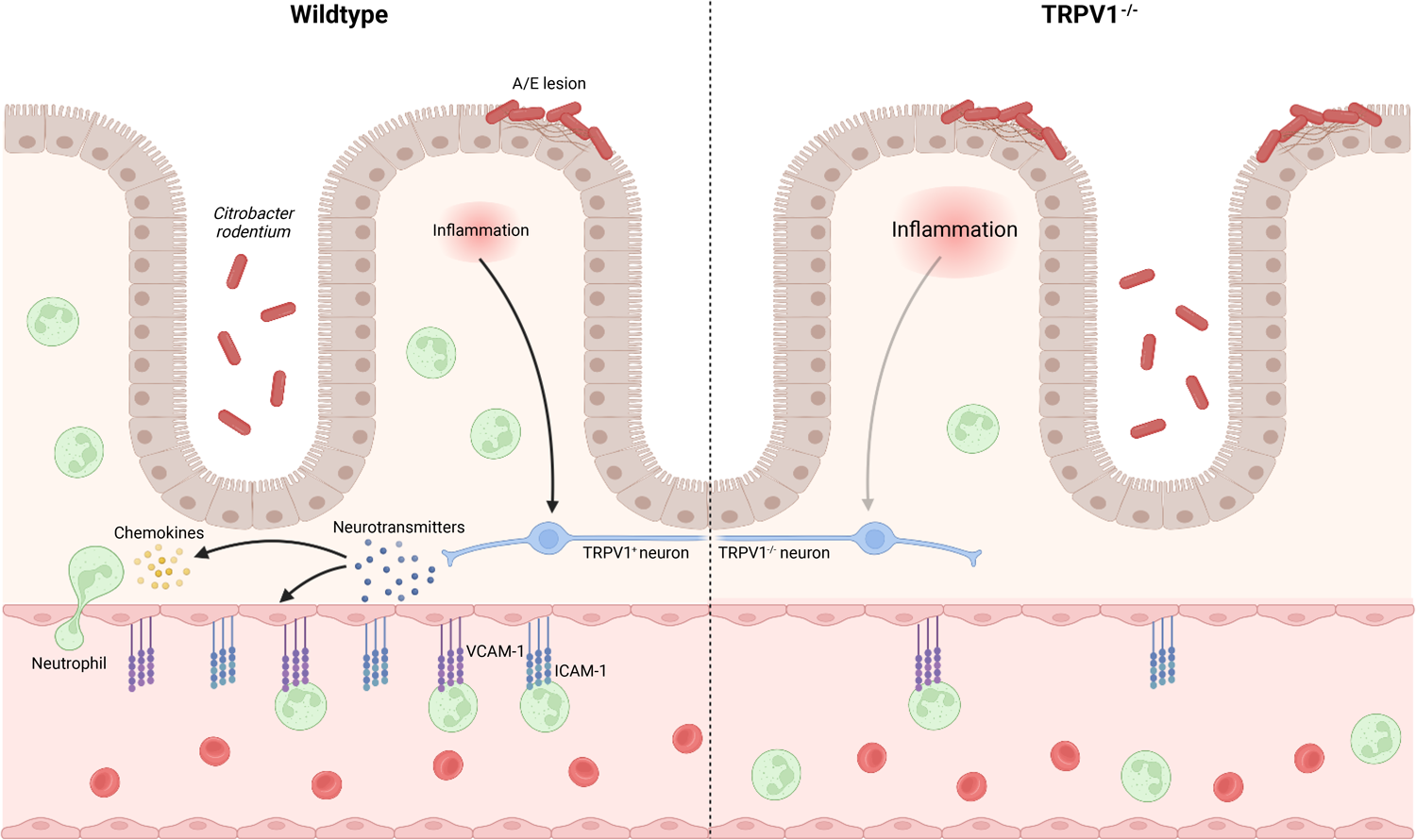
Neuronal TRPV1 signaling coordinates host protective immune responses during *C. rodentium* infection. TRPV1^+^ neurons promote the recruitment of neutrophils by upregulating cell adhesion molecules, ICAM-1 and VCAM-1, on blood endothelial cells and by promoting the release of the chemokine, CXCL6. These factors collectively promote extravasation of neutrophils from the blood vessel into the site of infection for clearance of *C. rodentium*. Mice lacking TRPV1 exhibit dramatic decreases in recruitment of neutrophils necessary for *C. rodentium* clearance from the colon.

## Discussion

Host immune responses to enteric bacterial pathogens are highly complex, requiring coordination between immune, epithelial, and stromal cells. Integral to these responses is the neuronal innervation of the intestinal tract, and the ability of these neurons to communicate with these various cell types through the release of specific neurotransmitters. Previously, we demonstrated a critical role for TRPV1^+^ sensory nociceptive neurons in the control of *C. rodentium* infection by ablation of these neurons; however, it was unclear if these host-protective effects were dependent on TRPV1 specifically, or by other receptors or secreted factors produced by these TRPV1^+^ neurons. Indeed deficiency in the related transient receptor potential cation channel, subfamily A, member 1 (TRPA1), which could also be expressed by these intestinal nociceptive neurons, significantly increased *C. rodentium* burden and tissue pathology (38). Using WT and TRPV1^-/-^ mice, we now demonstrate that this polymodal nociceptor is a critical component of the host response to enteric bacterial infection with *C. rodentium*.

Deficiency in TRPV1 not only increased bacterial burden in the feces and colon, but also exacerbated colonic histopathology. Intestinal epithelial cells have been reported to express TRPV1 (27, 28), and it has further been suggested that TRPV1 negatively regulate IEC proliferation (22). Although non-infected TRPV1^-/-^ mice exhibited increased crypt length compared to WT, the number of proliferating cells, indicated by Ki67 staining, was not different in uninfected mice. Moreover, epithelial cell organoid cultures where proliferation was measured by incorporation of the thymidine analog EdU was not different in WT or TRPV1^-/-^ derived cultures, and this rate of proliferation was not altered by treatment with the TRPV1 agonist capsaicin. Together these findings suggest that the increased crypt length and IEC proliferation observed in TRPV1^-/-^ compared to WT mice, was due to increased bacterial burden and the ensuing immune response, as opposed to altered IEC intrinsic TRPV1 signaling.

As a major component of the host response to *C. rodentium* infection depends on CD4^+^ T cells, we assessed the effect of TRPV1 deficiency in the recruitment of these cells. No significant difference in colonic T cells was observed in non-infected or *C. rodentium* infected TRPV1^-/-^ compared to WT mice. Quantification of the cytokines produced by colonic lamina propria T cells revealed only subtle but significantly reduced IL-17A production in infected TRPV1^-/-^ mice compared to WT. Previous reports found that during DSS induced colitis, mice with a mutation to constitutively activate TRPV1 showed a significant increase in CD4^+^ T cells producing IL-17A but not IFNγ (39). This was attributed to an increase in DC pro-inflammatory state, but here we show that, in a genetic knockout of TRPV1, IL-17A was significantly decreased in CD4^+^ T cells. These observations are in keeping with reduced expression of IL-17A in TRPV1^-/-^ T cells *in vitro,* although in contrast to that study we found no difference in IFNγ expression in CD4^+^ T cells *in vivo* (34). We also found that T cell culture demonstrated no difference in proliferation induced by TCR activation with or without co-stimulation, and no effect of the TRPV1 agonist capsaicin on proliferation in TRPV1^-/-^ compared to WT cells. These differences are most likely attributed to our analysis after *in vivo* infection compared to an *in vitro* assay with purified T cells. Although it is unclear what TRPV1 agonists would be present in an *in vitro* culture, these differences could also be attributed to unique ligands present during infection-induced inflammation *in vivo* compared to *in vitro*. In addition, since most *in vitro* assays utilize splenic T cells, it is unclear if expression of TRPV1 in CD4^+^ T cells is retained after recruitment to the lamina propria as an effector cell.

Although adaptive immune responses are critical for the clearance of *C. rodentium* infection and sterilizing immunity (32), there are critical and non-redundant roles for innate immune cells. These innate immune responses include bactericidal activity, phagocytosis, and release of chemokines and cytokines. During enteric bacterial infection, chemokine and cytokine production serve to increase the expression of host protective factors such as antimicrobial peptides and to increase blood endothelial cell adhesion molecules. Enteric infection resulted in significantly increased expression of cytokines including *Il6*, *Tnfα* and *Il1β*; however, at 10 days p.i., *Il6* and *Tnfα* were not different in TRPV1^-/-^ mice compared to WT while *Il1β* was significantly decreased in TRPV1^-/-^ mice, and at 29 days p.i., only *Il6* was significantly decreased in TRPV1^-/-^ mice compared to WT. While the expression of the anti-microbial protein *RegIIIγ* was significantly increased in infected TRPV1^-/-^ compared to WT mice, this is likely a response to the significantly increased bacterial burden in these animals. Neutrophils have been previously demonstrated to be indispensable antibacterial effectors during enteric bacterial infection.

Depletion, or genetic deficiency in chemokine receptors that dictate homing and migration of these cells into infected tissues, resulted in significantly increased bacterial burden and pathology (16). Our flow cytometric analysis revealed significantly reduced numbers of colonic neutrophils in *C. rodentium* infected TRPV1^-/-^ mice which is consistent with a role for TRPV1 in promoting host protective recruitment of these cells. Despite reduced recruitment of these cells, only *Cxcl6*, a neutrophil chemokine and antimicrobial peptide produced by activating resident macrophages and epithelial cells (35), was reduced in infected TRPV1^-/-^ mice. However, chemokines are well established to have compensatory roles, where deficiency in one chemokine is unlikely to result in a drastic phenotype (36, 37). Our analysis on the development of neutrophils from the granulocyte monocyte progenitor in the bone marrow of control and infected WT and TRPV1^-/-^ mice demonstrated no impact in development and maturation of these cells. These data indicate that any deficiency in the colonic tissue is therefore not simply due to a reduced ability to produce neutrophils in the absence of TRPV1. In support of this finding, prior studies with TRPV1^+^ neuronal ablation also showed no difference in the production of many immune cell types including neutrophils (40).

Recruitment of immune cells and extravasation into the infected tissue is dependent on the interaction of specific adhesion molecules on the immune and blood endothelial cells (41–44). Adhesion molecule expression and localization on the luminal surface of endothelial cells is well appreciated to increase during inflammation due to cytokines such as TNFα and IL-1β (45–47), prostaglandins (48), and bacterial products such as LPS (45, 49). Cytokines and products released during inflammation are not the only factors that can activate endothelial cells to promote recruitment of immune cells from the blood. Neurotransmitters released from sensory neurons such as SP and CGRP can act on blood endothelial cells increasing blood vessel permeability, and ICAM-1 on the endothelial cell surface (7, 50, 51). Our data demonstrate that colonic endothelial cells from *C. rodentium* infected TRPV1^-/-^ mice have significantly reduced ICAM-1 and VCAM-1 expression compared to WT mice. These findings agree with prior publications showing that the blockade of ICAM-1 attenuated colonic neutrophil recruitment during *C. rodentium* infection (41). Direct inflammatory effects of TRPV1^+^ neuronal activation have also been observed previously by selective activation of cutaneous sensory nociceptors using optogenetics to induce localized inflammation. Of particular interest, stimulation of these nerves induced IL-23 dependent recruitment of neutrophils and Th17 CD4^+^ T cells (52).

Neuronal TRPV1 dependent recruitment of neutrophils has also been shown during TLR agonist induced skin inflammation (40). In this study, the ablation of TRPV1 neurons had attenuated inflammation which was attributed to extravascular mechanisms; however, it is critical to note that adhesion molecule expression of leukocyte rolling was not performed during inflammation. In this context, our data suggests that TRPV1^+^ sensory neurons have a conserved ability to regulate inflammation in the intestinal tract. This ability of nociceptive neurons to control neutrophil recruitment in the intestinal tract is likely to have unique outcomes depending on the health of the host and the specific challenges that elicit this effect. In mouse models of colitis, neutrophils can exert host beneficial (53, 54) or detrimental effects (55, 56) depending on the model of disease. With these differential effects, it is tempting to speculate that this may be the basis for seemingly contradictory roles of nociceptive neuropeptides. As such, there appears to be considerable biological complexity, where sensory innervation, the signals released by these neurons, and the cells targeted may be capable of eliciting unique outcomes and suggests that, with further understanding, these processes could be used to enhance or reduce inflammation in a contextually meaningful manner. In a broader context, it has been shown that treatment-resistant IBD patients exhibited increased levels of neutrophils that correlated with increased expression of IL-1β and CXCL6 (57). Since TRPV1 has been implicated in other gastrointestinal disorders including IBD (58, 59), our data suggests a potentially broader context for conserved functions of TRPV1 in the gut that should be explored further.

---

**Fig. S1.**
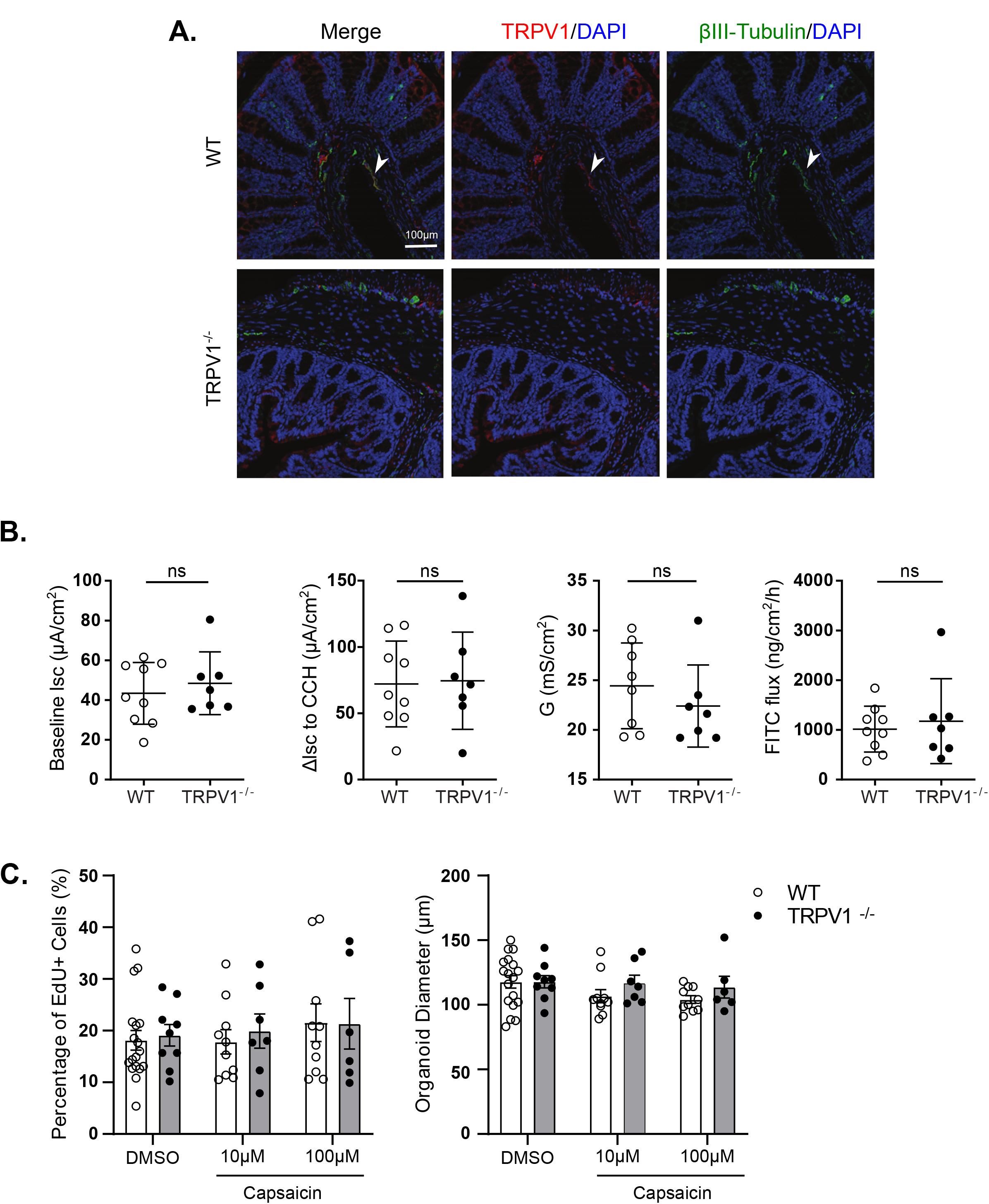
TRPV1 expressed in the colon does not affect gut permeability or epithelial cell proliferation. **(A)** Paraffin embedded colonic tissue of wild-type (WT) and TRPV1^-/-^ mice was stained with anti-TRPV1 (red), anti-βIII-tubulin (green), and DAPI (blue). **(B)** WT (open circles) and TRPV1^-/-^ (black circles) mice colons were assessed for gut permeability through Ussing chambers. Not significant. Student t test with 7-9 animals per group. **(C)** Proliferation of colon-derived organoids was assessed *in vitro* with EdU incorporation and diameter measurements. WT and TRPV1^-/-^ derived organoids were compared. Not significant. Student t test comparing WT to TRPV1^-/-^ with 7-11 organoids from 3 mice per group.

**Fig. S2.**
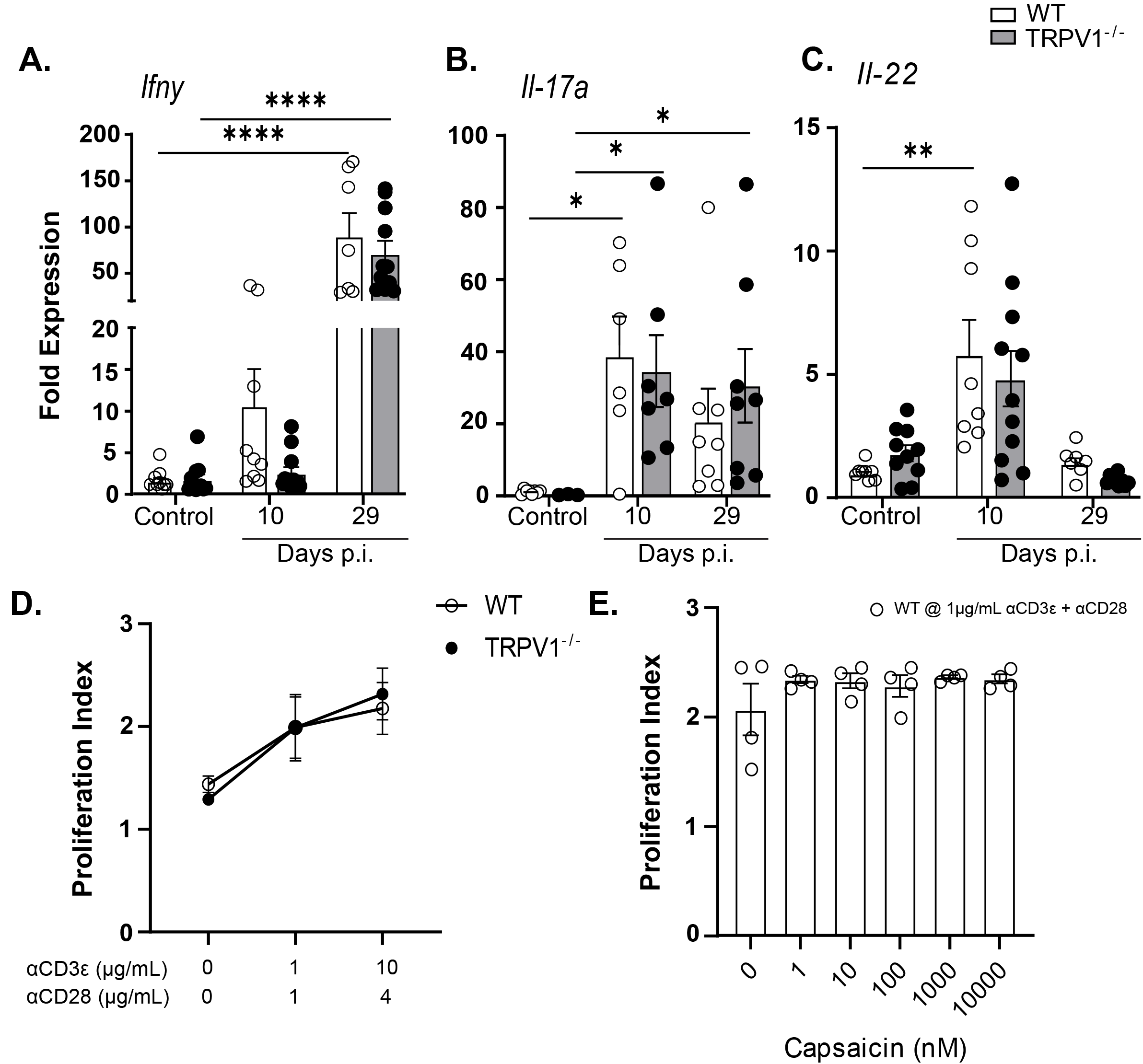
IFNγ, IL-17A, and IL-22 show no difference in mRNA transcripts at baseline and 10- and 29- days post-infection by *C. rodentium*. (A-C) Colonic tissue from wild-type (WT) and TRPV1^-/-^ mice was assessed by qPCR for expression of common T cell produced cytokines relevant to *C. rodentium* clearance such as (A) *Ifnγ*, (B) *Il17a*, and (C) *Il22*. Data are presented as mean ± standard error of the mean: *, P < 0.05, **, P < 0.01 and ***, P < 0.001; one-way ANOVA with post-hoc analysis using Tukey’s multiple comparisons test. 6–12 animals per group. **(D & E)** WT or TRPV1^-/-^ negatively selected CD3^+^ CD4^+^ T cells were stained with CellProliferation dye eFluor450 and cultured *in vitro* at described concentrations of anti-CD3ε and anti-CD28 antibodies for 72 hours and then analyzed by flow cytometry for proliferation index. **(D)** Comparison of WT and TRPV1^-/-^ T cell proliferation at different concentrations of anti- CD3ε and anti-CD28 antibodies. **(E)** WT T cells were cultured with 1 µg/mL of anti-CD3ε and 1 µg/mL of anti-CD28 and a dose response of the TRPV1 agonist capsaicin. After 72 hours, cells were analyzed by flow cytometry for proliferation index. Data are presented as mean ± standard error of the mean: not significant.

**Fig. S3.**
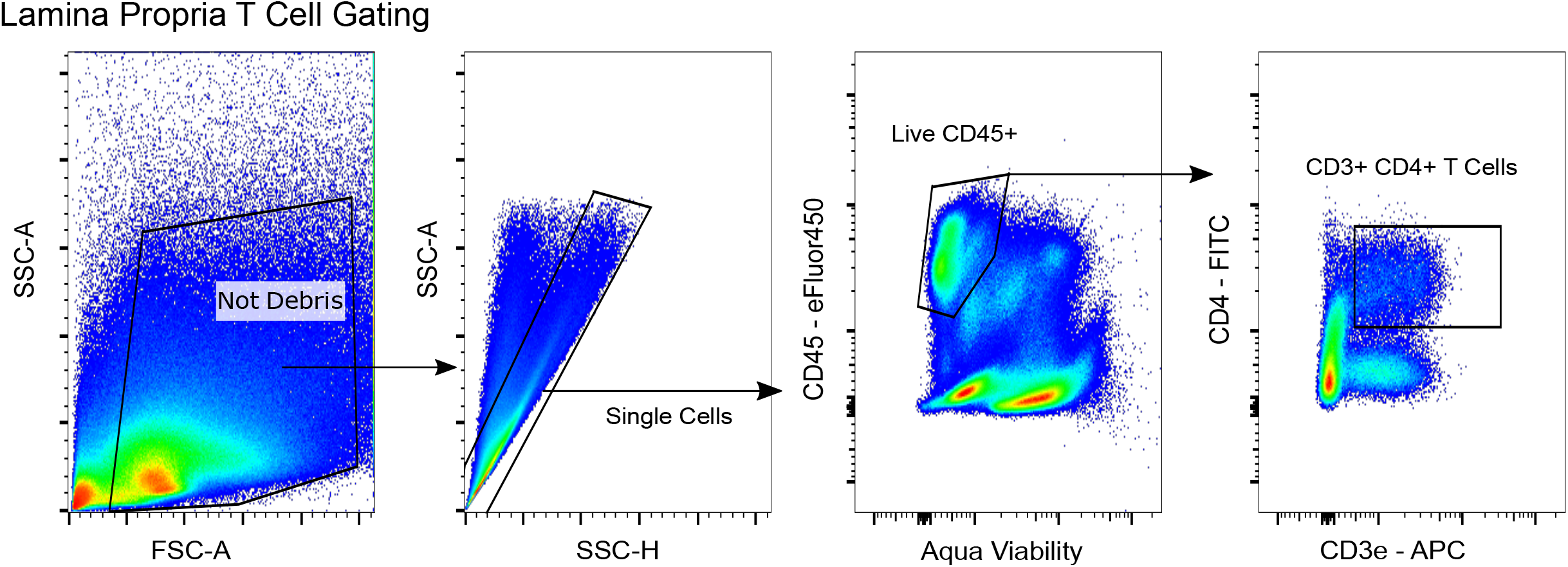
Lamina propria T cell gating strategy. Whole colon dissociated into a single cell suspension and stained for antibodies to identify live CD45^+^ CD3^+^ CD4^+^ T cells.

**Fig. S4.**
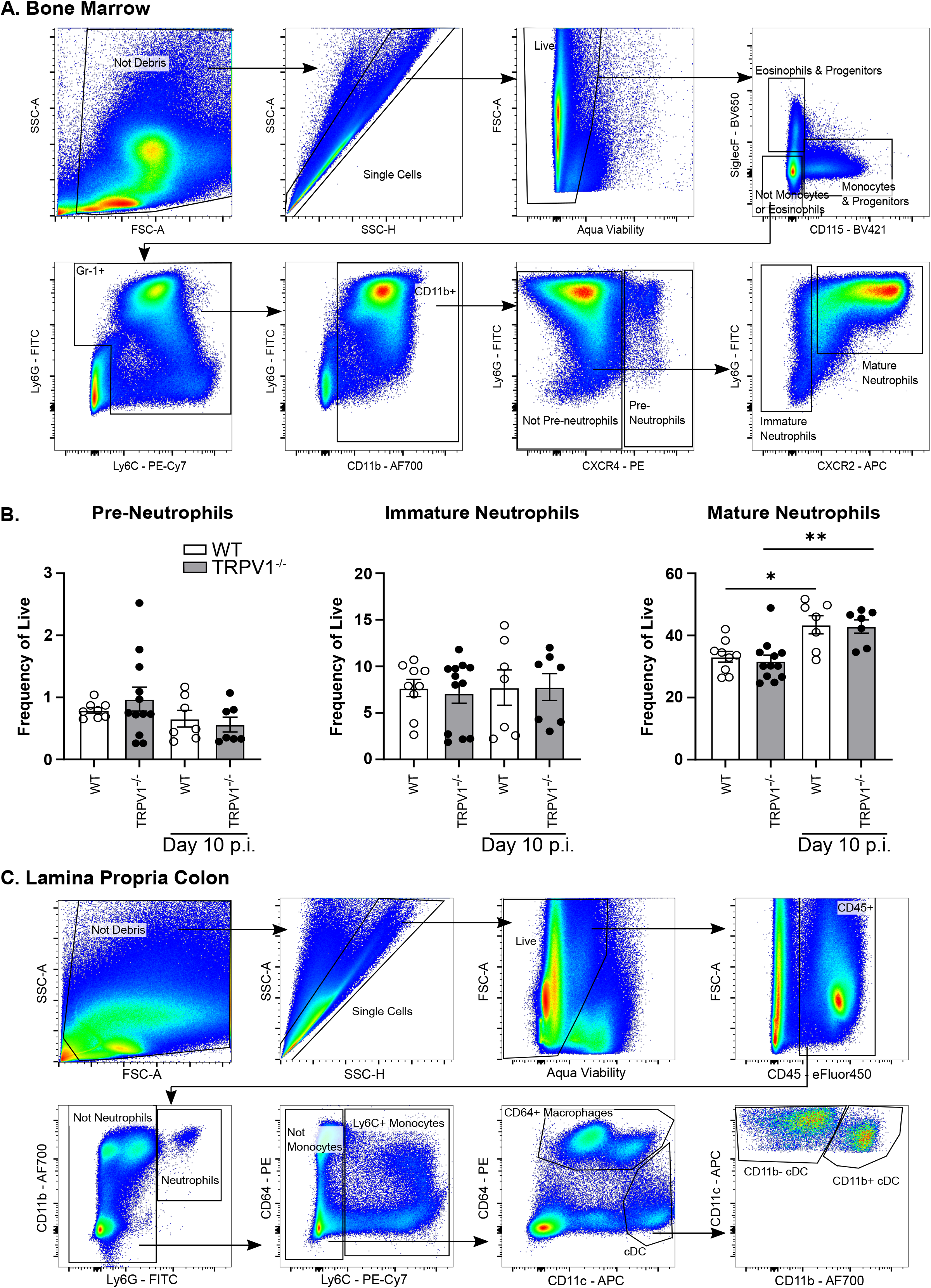
TRPV1 deletion did not alter neutrophil precursors in the bone marrow compartment. Wild-type (WT) and TRPV1^-/-^ mice had right femur bone marrow extracted at baseline and 10 days p.i. of *C. rodentium* and stained for pre-neutrophils (SiglecF^-^, CD115^-^, Gr- 1^+^, CD11b^+^, CXCR4^+^), immature neutrophils (SiglecF^-^, CD115^-^, Gr-1^+^, CD11b^+^, CXCR4^-^, CXCR2^-^), and mature neutrophils (SiglecF^-^, CD115^-^, Gr-1^+^, CD11b^+^, CXCR4^-^, CXCR2^+^, Ly6G^+^). **(A)** Bone marrow gating strategy and **(B)** frequency of live of each subset of neutrophil lineage shown. Data are presented as mean ± standard error of the mean: one-way ANOVA with Tukey post-test, with 7–12 animals per group, not significant. **(C)** Gating strategy for whole colon dissociated into a single cell suspension and stained for antibodies to identify live (CD45^+^, Ly6G^+^, CD11b^+^) neutrophils, (CD45^+^, Ly6G^-^, Ly6C^+^) monocytes, (CD45^+^, Ly6G^-^, Ly6C^-^, CD64^+^) macrophages, (CD45^+^, Ly6G^-^, Ly6C^-^, CD64^-^, CD11c^hi^) conventional dendritic cells (DC).

**Table S1.**
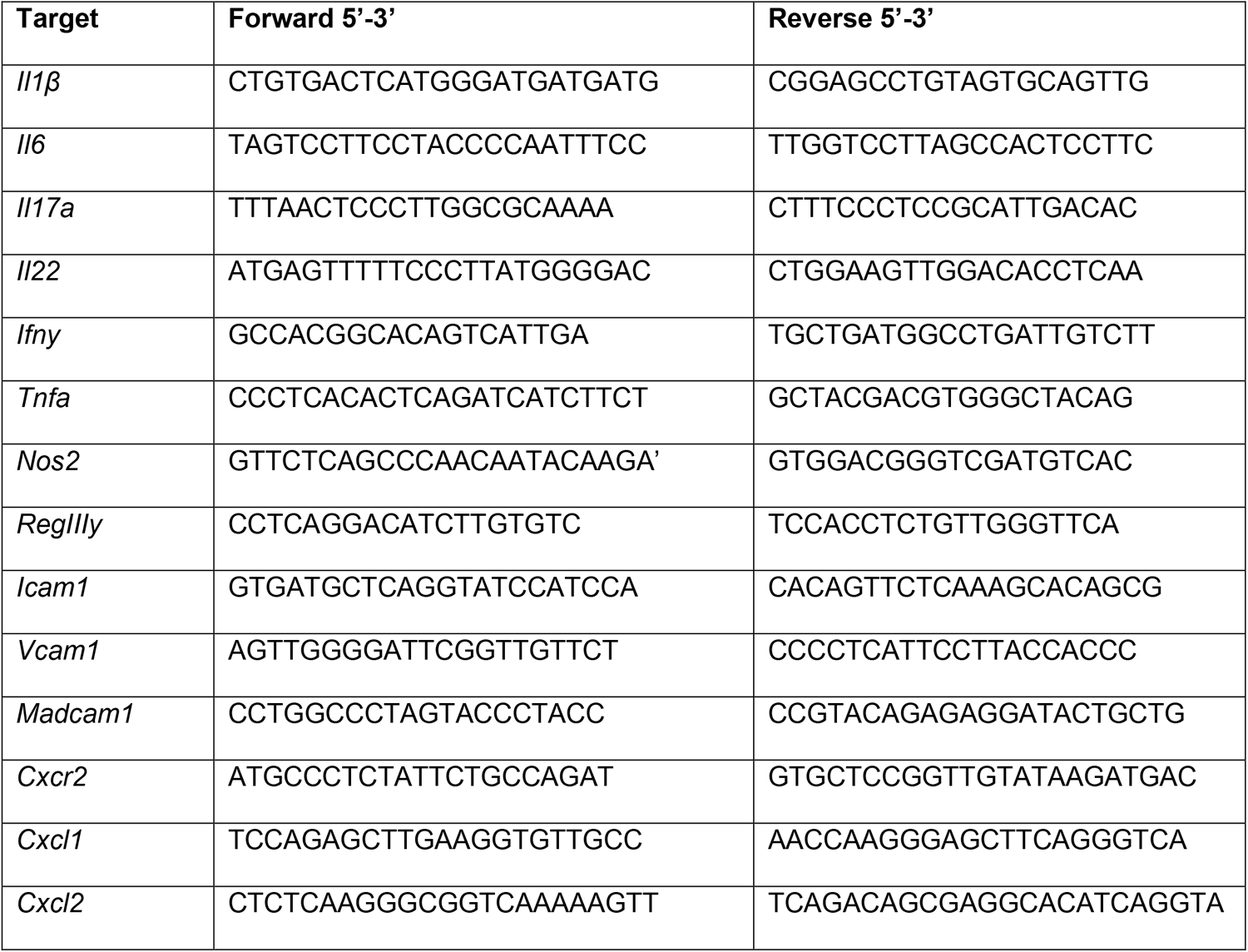
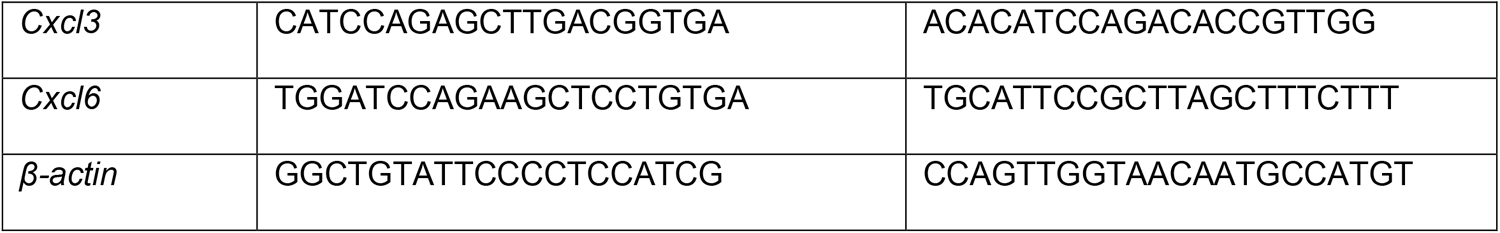
List of primers used for qPCR.

**Table S2.**
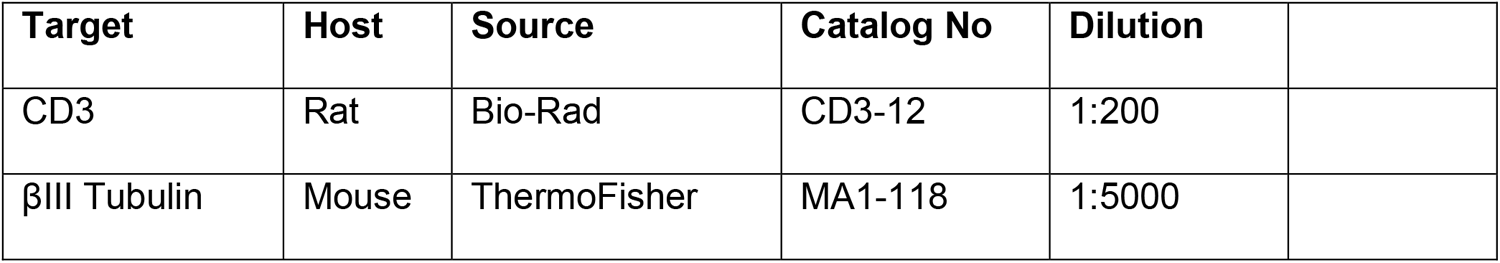
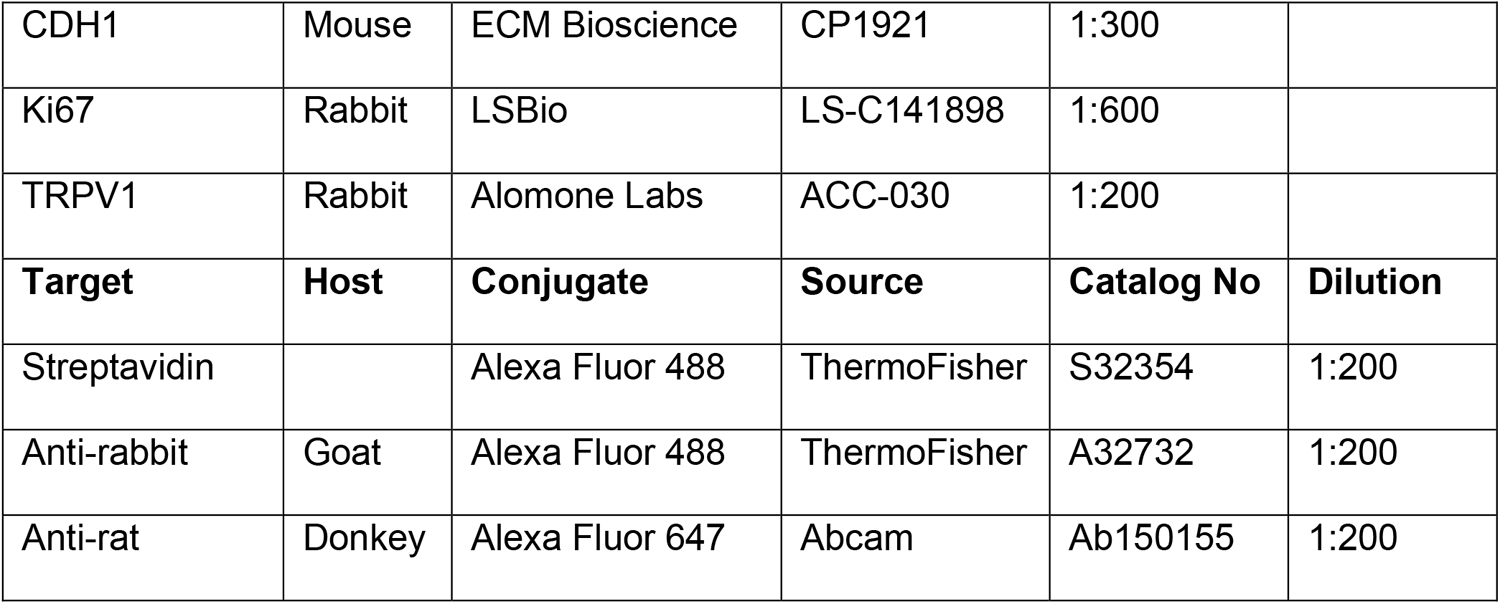
Antibodies used for Confocal Imaging.

| Target | Host | Source | Catalog No | Dilution |
| --- | --- | --- | --- | --- |
| CD3 | Rat | Bio-Rad | CD3-12 | 1:200 |
| βIII Tubulin | Mouse | ThermoFisher | MA1-118 | 1:5000 |

|  |  |  |  |  |  |
| --- | --- | --- | --- | --- | --- |
| CDH1 | Mouse | ECM Bioscience | CP1921 | 1:300 |  |
| Ki67 | Rabbit | LSBio | LS-C141898 | 1:600 |  |
| TRPV1 | Rabbit | Alomone Labs | ACC-030 | 1:200 |  |
| **Target** | **Host** | **Conjugate** | **Source** | **Catalog No** | **Dilution** |
| Streptavidin |  | Alexa Fluor 488 | ThermoFisher | S32354 | 1:200 |
| Anti-rabbit | Goat | Alexa Fluor 488 | ThermoFisher | A32732 | 1:200 |
| Anti-rat | Donkey | Alexa Fluor 647 | Abcam | Ab150155 | 1:200 |

**Table S3:** Flow cytometry antibodies.

| Antigen Target | Manufacture & Clone | Catalog number |
| --- | --- | --- |
| CD3 | Tonbo Biosciences 145-2C11 | 20-0031 |
| CD4 | BD Biosciences RM4-5 | 553047 |
| CD45 | Invitrogen 30-F11 | 48-0451-82 |
| Ly6G | Tonbo Biosciences 1A8 | 35-1276 |
| CD11b | Invitrogen M1/70 | 56-0112-82 |
| IFNγ | Invitrogen XMG1.2 | 25-7311-82 |
| IL-17A | BD Biosciences TC11-18H10 | 561020 |
| IL-22 | Invitrogen 1HBPWSR | 46-7221-82 |
| CD31 | Biolegend MEC13.3 | 102533 |
| gp38 | Biolegend 8.1.1 | 127405 |
| ICAM-1 | Biolegend YN1/1.7.4 | 116114 |
| VCAM-1 | Biolegend 429(MVCAM.A) | 105719 |
| MAdCAM-1 | Biolegend MECA-367 | 120710 |
| Fixable Live/Dead Aqua | ThermoFisher | L34957 |
| CD115 | BD Bioscience T38-320 | 743638 |
| SiglecF | BD Biosciences E50-2440 | 740557 |
| Ly6C | Invitrogen HK1.4 | 25-5932-82 |
| CXCR4 | Miltenyi Biotec REA107 | 130-118-682 |
| CXCR2 | Miltenyi Biotec REA942 | 130-115-635 |
| CD64 | Miltenyi Biotec REA286 | 130-118-684 |
| CD11c | Invitrogen N418 | 17-0114-81 |

